# Probabilistic clustering of cells using single-cell RNA-seq data

**DOI:** 10.1101/2023.12.12.571199

**Authors:** Joy Saha, Ridwanul Hasan Tanvir, Md. Abul Hassan Samee, Atif Rahman

## Abstract

Single-cell RNA sequencing is a modern technology for analyzing cellular heterogeneity. A key challenge is to cluster a heterogeneous sample of different cell types into multiple different homogeneous groups. Although there exist a number of clustering methods, they do not perform well consistently across various datasets. Moreover, most of them are not based on probabilistic approaches making it difficult to assess uncertainties in their results. Therefore, in spite of having large cell atlases, it is often quite difficult to map cells to types. In addition, many of the methods require prior knowledge such as marker gene information for each type. Also due to technological limitations, dropouts of gene expressions may occur in the data which is not taken into account in other methods. Here we present a probabilistic method named CellHorizon for clustering scRNA-seq data that is based on a generative model, handles dropouts and works without any prior marker gene information. Experiments reveal that our method outperforms current state-of-the-art methods overall on six gold standard datasets.

## Introduction

Single-cell RNA sequencing (scRNA-seq) technology has become a popular tool to understand the relationship between cells and genes [1]. Even with its growing usage, scRNA-seq data analysis remains a difficult task [24]. There is substantial variability in the data. In addition, the gene expression measurements in the scRNA-seq data are low and sparse, with many “false” zero count observations known as dropout events [17]. The dropout event occurs because of the low RNA capture rate and the low sequencing depth per cell.

Another challenge is that in most scRNA-seq studies, cell types are not known with certainty. Researchers typically use unsupervised clustering techniques to organize cells into sets. Cell types can be identified based on the clustering results [12]. Jaitin *et al*. showed that dissection of tissues into mixtures of cell types is a difficult task [10]. It is a standard procedure to use marker genes after cells are grouped into clusters. Ringeling *et al*. observed that Seurat [5] fails to distinguish CD4 T cells and CD8 T cells based on transcriptomic data alone, Hence, the authors used both marker genes and protein marker expressions (ADT, antibody-derived tags) to estimate cell types. However, they were still unable to label one-fifth of the cells [19].

Unsupervised methods like Leiden [23], Louvain [4] and SC3 [13] are some of the most popular clustering methods. Leiden [23] algorithm, an improved version of Louvain [4] moves its nodes locally to optimize a modularity function. Then it refines the partition and creates an aggregate network. These are computationally very expensive. Another unsupervised clustering method SC3 [13] generates distance matrices like Euclidean, Pearson, and Spearman, and then uses their PCA transformation to get their eigenvectors. Clustering is done on each of these eigenvectors and their consensus is taken as the result. This method requires fixing a large number of parameters upon which the quality of the clustering depends. Most of these parameter values can not be selected intuitively. An autoencoder-based method scGAC [7] takes the raw-count matrix as input and after normalizing and log transformation it gets the expression matrix. Using auxiliary matrices and PCA, it generates a feature matrix that is optimized by Graph Attentional Autoencoder (GAA). Based on the latent space representation of the GAA, it optimizes a membership matrix based on k-means clustering to assign each cell to one specific cluster. EDClust [26] is a statistical probabilistic clustering method that focuses on subject-specific factors of single-cell data. It uses the Dirichlet-multinomial mixture model to represent expression counts in multi-subject scRNA-seq data. It follows an EM–MM hybrid method. In the E-step, it calculates a posterior probability for cluster assignment and in M-step, it optimizes a loss function. Then to incorporate cell type-specific and subject-specific effects it uses a MM algorithm.

These unsupervised algorithms often use a low dimensional representation like PCA of count data and cluster the cells based on this. The low-dimensional representations measure linear similarity among cells but fail to capture the latent relationship which leads to sub-optimal clustering [9].

In addition to the clustering methods, there exist a number of supervised and semi-supervised methods like CellAssign [29] and SCINA [31]. Both CellAssign [29] and SCINA [31] require prior information. CellAssign [29] is a probabilistic approach to assign cells to types. Its main focus is to automate the process of annotation rather than clustering cells. For this, it requires additional marker gene information. However, determining the marker genes of a cell type that is not well known is a challenging task. CellAssign [29] also does not handle dropouts which is a key factor in single-cell clustering. Tabula Muris [22], which contained scRNA-seq data for about 120,000 cells from 20 organs and tissue types in mice had a dropout rate of up to 93% [18].

In this work, we propose an unsupervised probabilistic clustering method based on CellAssign [29] that does not require any prior marker gene information and models the expression data using negative binomial distribution. As it is a probabilistic approach it performs well when cells may have characteristics that span multiple clusters and helps us to understand overlapping relationships between clusters. It allows us to capture the uncertainty associated with each cell’s assignment to a cluster. It also takes dropout into account by associating a dropout rate with each gene so that, dropout and actual zero value in the expression can be differentiated. Our method also does not require any additional parameter value and it is based on expression count data rather than any low-dimensional representation of it.

## Method

In scRNA-seq clustering, given an expression matrix (*n × g*) of *n* cells and *g* genes, the challenge is to cluster the cells into *c* different clusters. Our method achieves this through a generative probabilistic approach and using negative binomial distributions to represent the count data. At first, the mean and dispersion parameters of the negative binomial distributions are initialized from the expression matrix. Then an EM algorithm is used to iteratively re-estimate the mean, dispersion and other parameters of the model. The EM algorithm terminates when the model converges. Then the probability for each cell being each type is calculated from the corresponding negative binomial distributions.

### Handling Dropouts

One of the key focuses of our method is handling dropouts. ZingeR [2] is a tool to model dropouts. It handles dropouts using a two component mixture model. The dropout is modeled by a point mass at zero and the original count as a negative binomial distribution. Then the probability that a zero expression count belongs to the count component is the negative binomial component divided by the mixture component. Our method uses a dropout parameter *d*_*g*_ for each gene which is the probability of dropout for a gene *g*.

### Generative Model

In the generative model of our algorithm, *z* or cell type indicator is determined by the prior probabilities *π*_*c*_. Based on *z*, we get the parameters of our negative binomial distribution *μ* and *ϕ* for each gene. Based on this negative binomial distribution, we sample the initial raw count value *y*_1_. Again *d*_*g*_ or dropout rate is used to determine *w* which is the probability of count value coming from zero distribution. *y* is the final count value that is determined by *y*_1_ and this incorporated *w*.

### Initialization

For each gene, we calculate the mean and variance column-wise. Then using Equation 1, we calculate the dispersion *ϕ*.

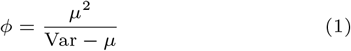

where *μ* is the mean and Var is the variance. So, *μ* and *ϕ* parameters for negative binomial distribution are initialized.

### EM Algorithm

The EM algorithm [15] or expectation–maximization algorithm is an iterative method of maximizing the likelihood of a statistical model by varying parameter estimation. The E step or expectation step determines an expectation of the log-likelihood of the model using the current estimates of the parameters. The M step or maximization step re-estimates the parameters using a loss function. This two steps are performed iteratively until a convergence criterion is met.

### E Step

In the E step, at first, we define two boolean variables *λ*_*ng*_ and *τ*_*ng*_ as follows:

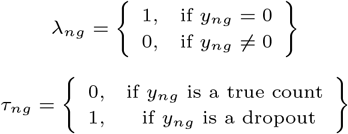

We calculate the probability that a zero value in the gene expression came from dropout *w*_*ng*_ as follows:

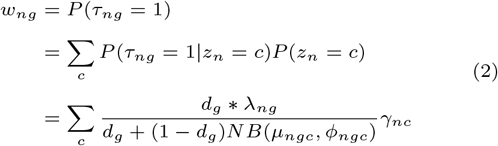

Then we calculate the mean *μ*_*ngc*_ of the negative binomial distribution using the following equation:

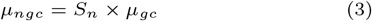

where *S*_*n*_ is the size factor of cell *n*. Using the mean *μ*_*ngc*_ and dispersion *ϕ*_*ngc*_ we calculate the probability that *n*^*th*^ cell is of type *c*.

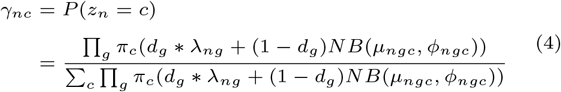

### M-step

In the M-step, we minimize the loss function

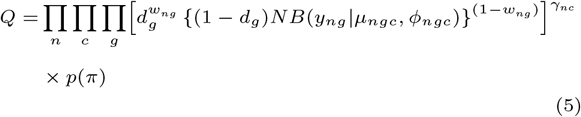

or equivalently its logarithm

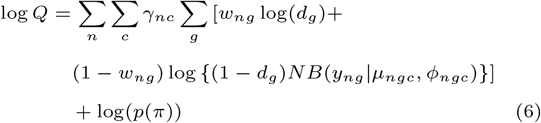

using Adam optimizer. The loss function also has two parts: one coming from negative binomial distribution and another from dropout. We update the parameters *d*_*g*_, *μ*_*ngc*_, *ϕ*_*ngc*_, *π*_*c*_ which are used for the next iteration of EM.

### Convergence Criteria

We use two types of convergence criteria for the EM algorithm. The model is assumed to be converged if any of these conditions are met.

- **Max Epoch:** The EM algorithm runs at most 400 epochs, then it stops and returns the results.
- **Early Stopping:** The model is stopped early and returned if for 15 consecutive iterations, the loss does not improve.

### Adam Optimizer

Adam optimizer [11] is used in the M step of our algorithm. It is a stochastic gradient descent method and it uses the adaptive estimation of first-order and second-order moments to minimize a loss function. In our method Equation 6 is minimized by Adam optimizer by changing the values of *d*_*g*_, *μ*_*ngc*_, *ϕ*_*ngc*_, and *π*_*c*_.

## Results

### Dataset

The input for our clustering method is a gene expression matrix where each row corresponds to a cell and each column represents a gene. We compared our model with other state-of-the-art models using six gold standard datasets, six silver standard datasets, and one simulated dataset.

#### Simulated Dataset

We generated a simulated dataset of 5 cell types of 9156 cells and 24 genes. We randomly selected 6 to 12 marker genes for each type of cell. For each gene and each cell type one negative binomial distribution was taken and used to sample the expression count. The mean *μ* for marker genes was taken two times more than the non-marker genes.

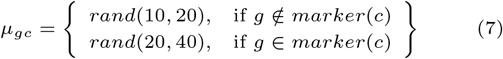

To calculate the dispersion parameter *ϕ*, we randomly generated variance Var.

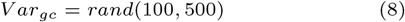

Then we can calculate *ϕ* using the following equation:

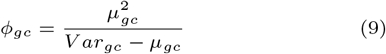

Dropout rate *d*_*g*_ was chosen as follows:

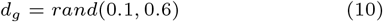

We assumed the size factor was 1 for all cells. Then using *μ*_*gc*_ and *ϕ*_*gc*_, negative binomial distributions were created and count values *y*_*ng*_ were sampled from them. Dropout rate *d*_*g*_ was used to incorporate the effect of dropout in our simulated dataset.

#### Gold-standard Datasets

Six gold-standard datasets and six silver-standard datasets were used for our comparison. Goolam, Kolodziejczyk, Yan, Deng, Pollen, and Biase are considered as gold-standard datasets because in these datasets the cell sub-populations were chosen from distinct biological stages or conditions [25].

#### Silver-standard Datasets

In addition, we analyze Worm neuron, Human kidney, 10x pbmc, Mouse retina raw, CITE CBMC and TAM FACS datasets. They are considered silver-standard datasets because in these the cells were labeled according to the authors’ computational analysis and biological knowledge [25].

### Preprocessing

We performed four preprocessing steps in our algorithm to filter the data and determine the genes whose expressions are highly differentiable across different cell types.

- Gene filter: We included genes that are expressed in at least three cells.
- Cell filter: A minimum of 200 genes expressed was required for a cell to pass filtering.
- We selected 1% most highly variable genes (HVGs) that have the highest variance across all cells. They have the greatest potential to distinguish between different cell types.
- Finally we applied log normalization to account for high variance in expression count.

### Performance Evaluation Metrics

We used Adjusted Rand Index (ARI) an improved version of Rand Index (RI) to compare the performance of our method with other methods using true and predicted cell types. ARI computes the similarity between two clusters on a scale from 0 to 1. ARI scores are higher if the predicted clustering is close to the true clustering.

### Result on Simulated dataset

First, we analyze the simulated dataset to assess how well our method can determine the parameters of our model: mean, dispersion and dropout rate.

From Fig 3 and Fig 4, we find that the predicted mean and dropout are similar to the original mean and dropout. Also, we can observe that the range of the original mean and dropout that is 10-40 and 0.1-0.6 are equal to the range of predicted mean and dropout of 10-40, and 0.1-0.6 respectively. We also find that our method has a very high ARI score of 0.948 in the simulated dataset. We varied the mean and dropout rate of the simulated dataset and found that our method could perform consistently with different ranges of mean, and dropout values. Hence, our method could determine the parameters of the negative binomial distributions precisely and cluster the cells accurately in the simulated dataset.

**Fig. 1.**
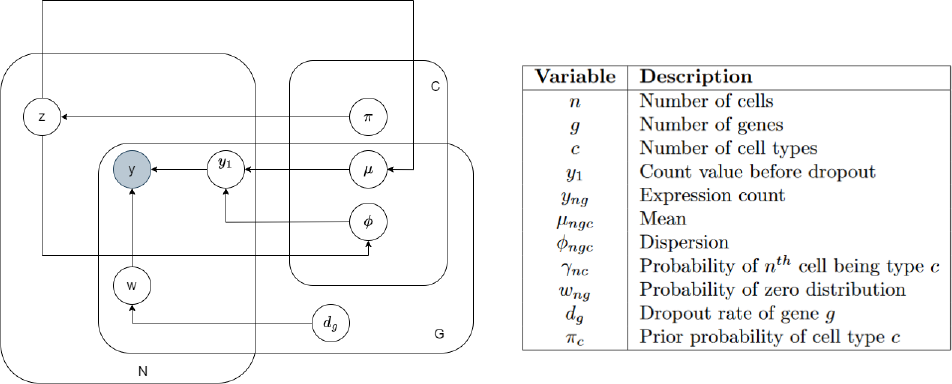
Generative Model

**Fig. 2.**
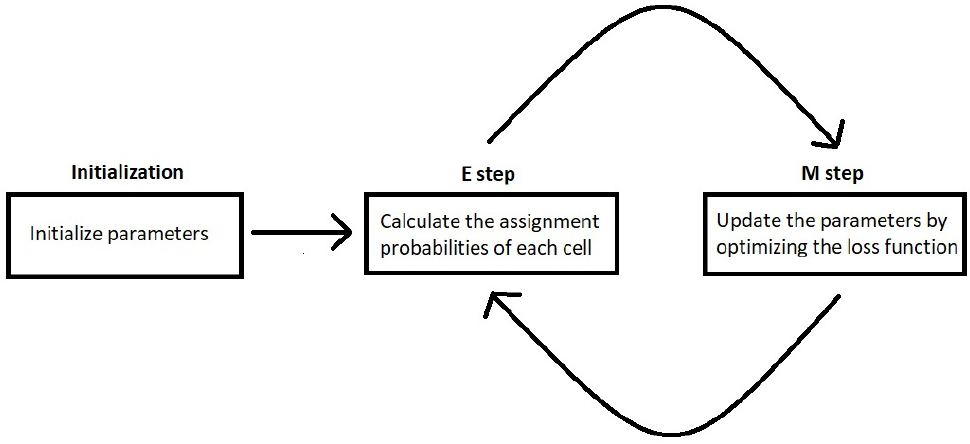
EM Algorithm

**Fig. 3.**
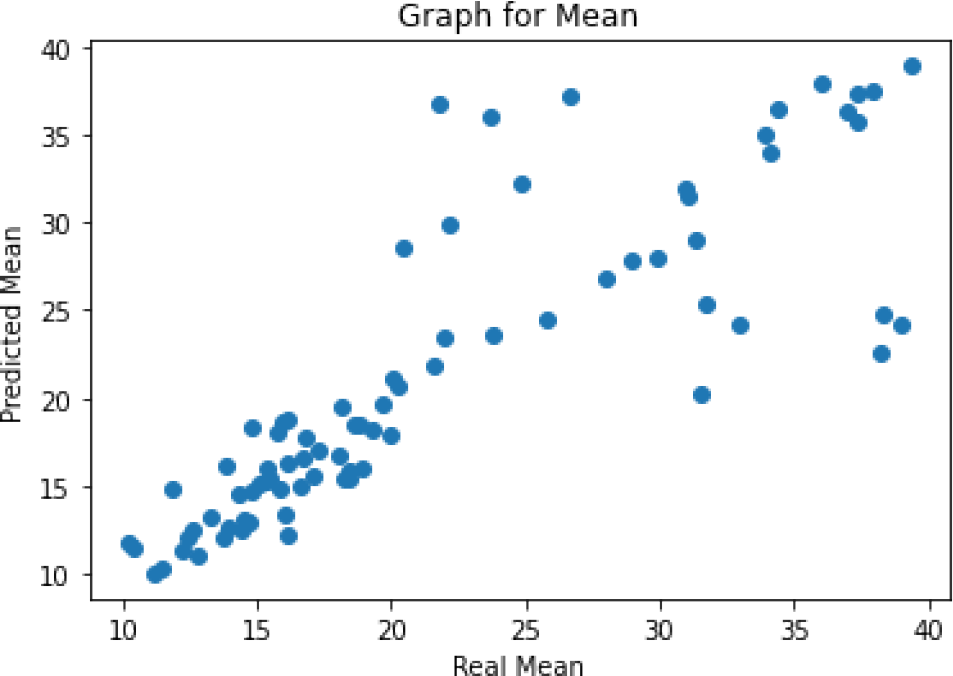
Real vs Predicted Mean

**Fig. 4.**
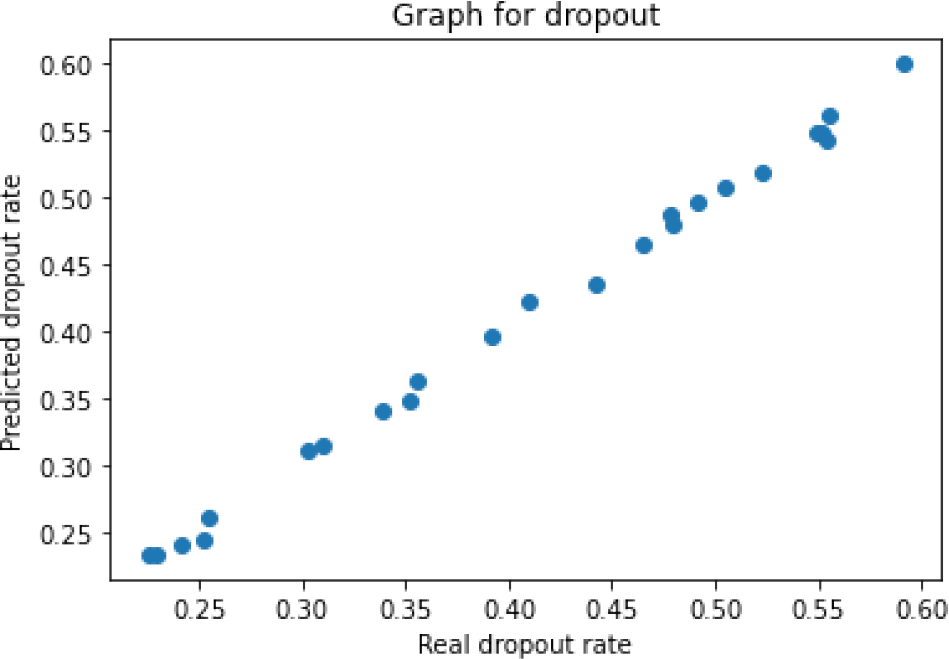
Real vs Predicted Dropout

**Fig. 5.**
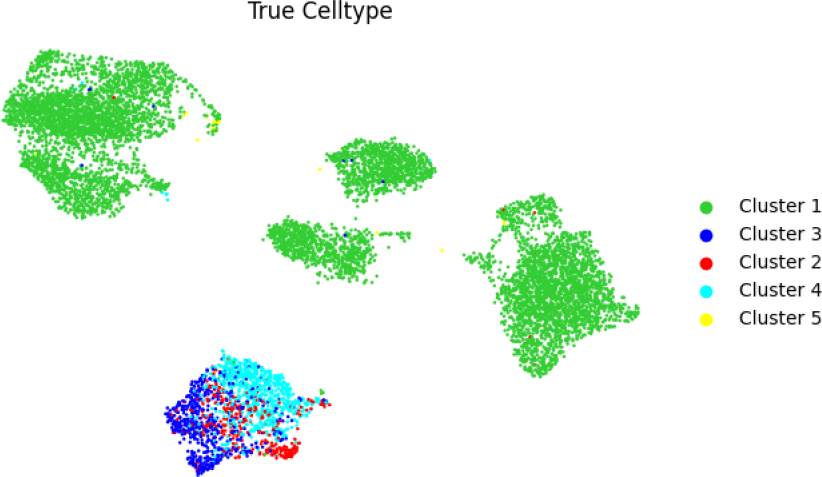
True celltype

**Fig. 6.**
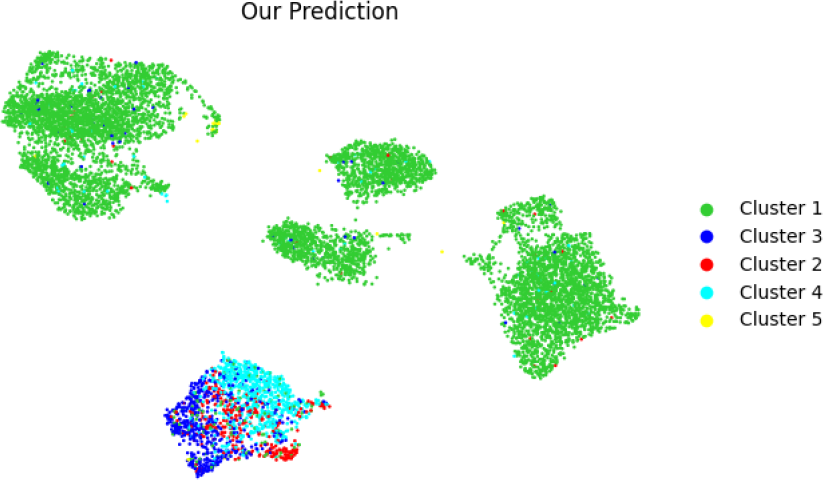
Predicted celltype

**Fig. 7.**
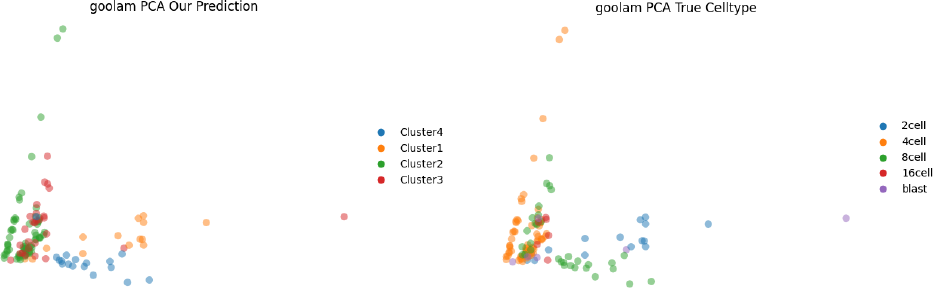
Goolam pca

**Fig. 8.**
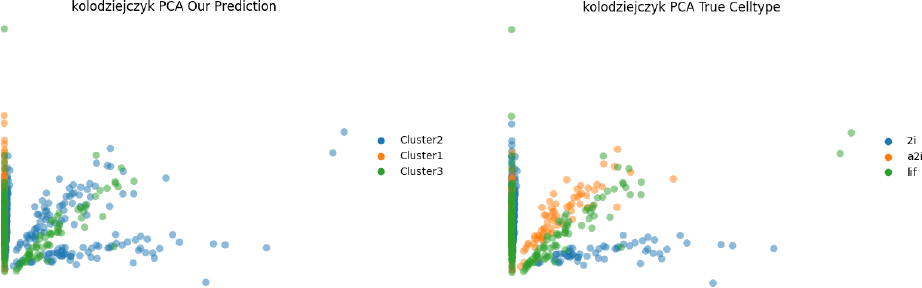
Kolodziejczyk pca

**Fig. 9.**
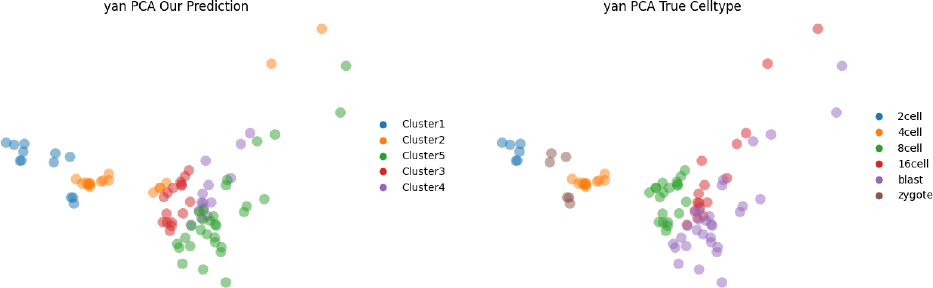
Yan pca

**Fig. 10.**
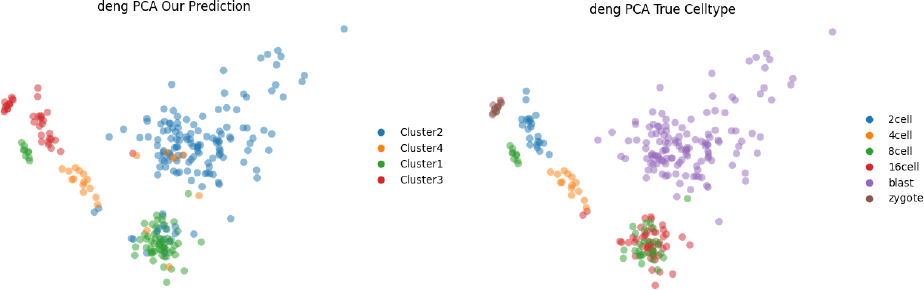
Deng pca

### Result on Real dataset

Next we analyze the six gold standard and six silver standard real datasets and compared the ARI of 5 state-of-the-art clustering methods with our method.

Table 3 shows the ARI of the methods on the gold standard datasets. As the cell sub-populations were chosen from distinct biological stages or conditions in gold datasets, they are more reliable than silver datasets where the labels were given using authors’ computational analysis. Our method outperforms all the other methods on average in 6 gold datasets. In the Goolam and Deng datasets, our method has a significantly higher ARI score than the second highest performing methods Leiden and scGAC respectively. Our method showed poor performance in only the Biase dataset. A possible reason is the number of cells is small in this dataset which may lead to inaccurate estimation of parameters.

**Table 1.** Gold-standard Datasets.

| Dataset Name | Total Cells | Total Genes | Cell Types |
| --- | --- | --- | --- |
| Goolam [9] | 124 | 41428 | 5 |
| Kolodziejczyk [14] | 704 | 38616 | 3 |
| Yan [27] | 90 | 20214 | 6 |
| Deng [8] | 268 | 22431 | 6 |
| Pollen [16] | 301 | 23730 | 11 |
| Biase [3] | 49 | 25737 | 3 |

**Table 2.** Silver-standard Datasets.

| Dataset Name | Total Cells | Total Genes | Cell Types |
| --- | --- | --- | --- |
| Worm Neuron [6] | 4186 | 13488 | 10 |
| Human Kidney [28] | 5685 | 25215 | 11 |
| 10x PBMC [32] | 4271 | 16653 | 8 |
| Mouse Retina [20] | 14653 | 11422 | 19 |
| CITE CBMC [21] | 8617 | 2000 | 15 |
| TAM FACS [30] | 110824 | 22966 | 126 |

**Table 3.**
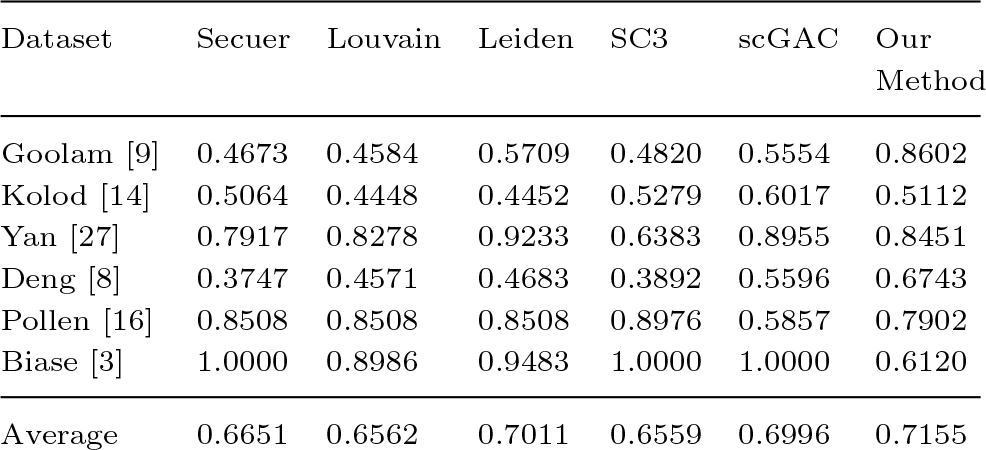
Gold-standard Datasets ARI Comparison.

In the case of silver datasets Securer, Louvain and scGAC (Tab 4) failed to run TAM FACS due to very high memory demands. Moreover, scGAC could not run Mouse Retina.

#### Running time

Table 5 shows running times of the methods. We find that our method is the third fastest method overall. The two methods faster than our method are SC3 and Secuer. However, their ARI scores are less than our method. Hence, no other method performed better in less time than our method.

**Table 4.**
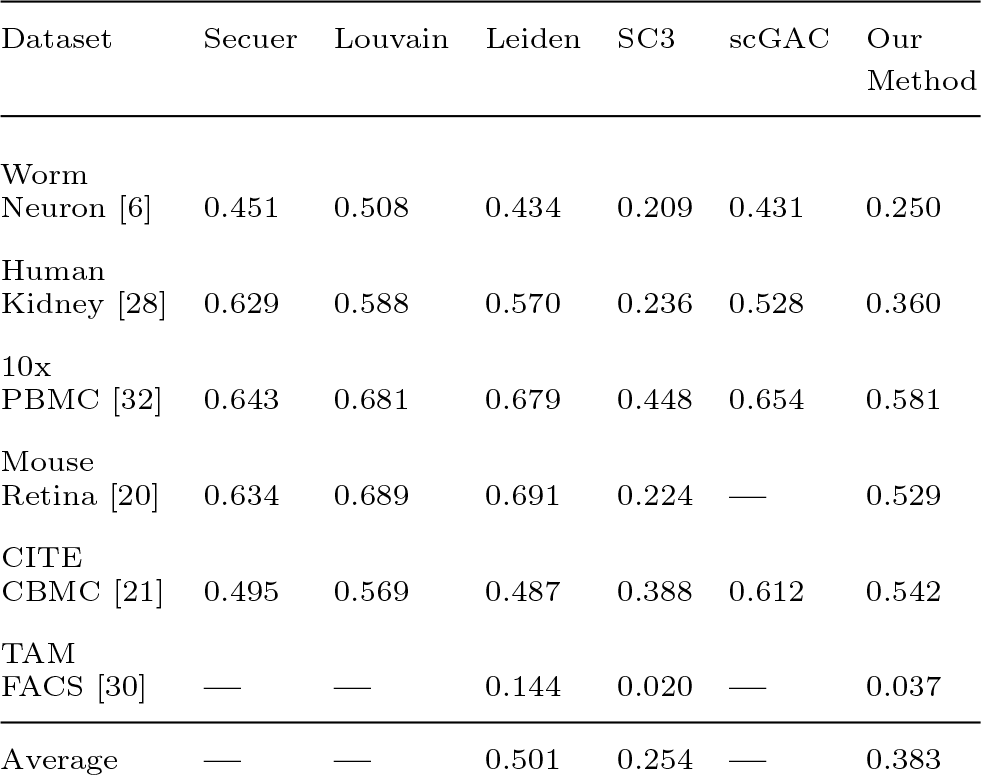
Silver-standard Datasets ARI Comparison.

**Table 5.** Gold-standard Datasets Time Comparison in milliseconds.

| Dataset | Secuer | Louvain | Leiden | SC3 | scGAC | Our Method |
| --- | --- | --- | --- | --- | --- | --- |
| Goolam [9] | 2849 | 23869 | 18972 | 1117 | 39492 | 11265 |
| Kolod [14] | 7284 | 24419 | 13336 | 4941 | 136261 | 12962 |
| Yan [27] | 3968 | 23779 | 12347 | 771 | 33388 | 10776 |
| Deng [8] | 5712 | 25895 | 12238 | 2294 | 41409 | 11407 |
| Pollen [16] | 4108 | 24800 | 11932 | 2400 | 44244 | 12307 |
| Biase [3] | 3836 | 22516 | 12440 | 666 | 28768 | 11513 |
| Average | 4626 | 24213 | 13544 | 2031 | 53927 | 11705 |

## Discussion

We generated the PCA visualization of the Gold datasets and analyzed the discrepancy between our clustering and ground truth celltypes visually. Moreover, we determined the probable marker gene list using both true celltypes and our predicted celltypes using scanpy. We calculated the number of common marker genes between these two lists for all possible combination of true celltypes and predicted celltypes.

### Goolam dataset

We can see in the true celltype, the 8 cell cluster is separated into two parts. But in our prediction, those two parts are clustered into different groups and also in the PCA visualization they appear to belong to different clusters. Moreover, there is a celltype blast in the true celltype which is a cluster of very few cells scattered around. As few scattered cells rarely make any cluster, our method did not consider it as a separate cluster. In further analysis, we found that blast has the lowest or second lowest number of common marker genes with all predicted clusters. This is possibly why blast was not considered as a different cluster by our method.

### Kolodziejczyk dataset

Our method merges some a2i cells into 2i cells. This is due to the fact that between 2i and a2i cells, the Spearman correlation coefficient of mean gene expression levels is very high (0.95). It is also reflected in the common marker list. For lif celltype, cluster 3 has the highest number of common marker genes much more than other clusters but for 2i and a2i both cluster 1 and cluster 2 have roughly the same number of common marker genes.

### Yan dataset

We can see that in true celltype the zygote cluster is divided into two parts and another cluster 4 cell is placed between them. Our prediction combines these two parts with cluster 1.

### Deng dataset

In the Deng dataset, we see that in true celltype the 16 cells and 8 cells are overlapping with each other. Our prediction does not have any overlapping and considers all of them as cluster 1.

### Biase dataset

Biase is a small dataset containing only 49 cells. Our ARI was the lowest in the Biase dataset. It is due to the fact that fitting a distribution with such small number of samples is relatively hard.

## Conclusion

Here we have presented a probabilistic unsupervised clustering algorithm for single-cell RNA-seq data based on a generative model. It models the expression data using negative binomial distributions. The method does not use any prior marker gene information. We also consider and tackle the effect of dropout to get a better quality clustering. We tested our method on the simulated dataset to evaluate the accuracy of parameter estimation and find that it can accurately estimate the means and dropout rates of genes. We then compared our method with other methods on Gold and Silver datasets. It was shown that our method is consistent and overall outperforms other methods in the gold datasets.

In future, a number of directions may be explored. We have used our method only for detecting clusters. But we can easily modify it to track the progress of cell development. Also, in the simulated dataset we have successfully estimated the mean and dropout parameters. However. there is scope for improvement in the estimation of the dispersion parameter. The method may enable finding the characteristics of unknown cell types from a heterogeneous cell population. So we may be able to detect new types of cells as well as subclusters.

Again, in tumor and cancer cells, not all the cells are malignant. There may be healthy cells beside so we would use the clustering algorithm to detect outliers. Doublet detection can be another promising aspect of our method as it can use the advantage of probabilistic assignment.

## Funding

This work was partially supported by the Research and Innovation Centre for Science and Engineering (RISE), Bangladesh University of Engineering and Technology (BUET).

## Notes

### Competing Interest Statement

The authors have declared no competing interest.

## References

1. Rhonda Bacher and Christina Kendziorski. Design and computational analysis of single-cell rna-sequencing experiments. Genome biology, 17(1):1–14, 2016.

2. K Berge Van den, Charlotte Soneson, and ML Love. zinger: unlocking rna-seq tools for zero-inflation and single cell applications. bioRxiv, 2017.

3. Fernando H Biase, Xiaoyi Cao, and Sheng Zhong. Cell fate inclination within 2-cell and 4-cell mouse embryos revealed by single-cell RNA sequencing. Genome Res., 24(11):1787–1796, November 2014.

4. Vincent D Blondel, Jean-Loup Guillaume, Renaud Lambiotte, and Etienne Lefebvre. Fast unfolding of communities in large networks. Journal of statistical mechanics: theory and experiment, 2008(10):P10008, 2008.

5. Andrew Butler, Paul Hoffman, Peter Smibert, Efthymia Papalexi, and Rahul Satija. Integrating single-cell transcriptomic data across different conditions, technologies, and species. Nature biotechnology, 36(5):411–420, 2018.

6. Junyue Cao, Jonathan S Packer, Vijay Ramani, Darren A Cusanovich, Chau Huynh, Riza Daza, Xiaojie Qiu, Choli Lee, Scott N Furlan, Frank J Steemers, Andrew Adey, Robert H Waterston, Cole Trapnell, and Jay Shendure. Comprehensive single-cell transcriptional profiling of a multicellular organism. Science, 357(6352):661–667, August 2017.

7. Yi Cheng and Xiuli Ma. scgac: a graph attentional architecture for clustering single-cell rna-seq data. Bioinformatics, 38(8):2187–2193, 2022.

8. Qiaolin Deng, Daniel Ramsköld, Björn Reinius, and Rickard Sandberg. Single-cell RNA-seq reveals dynamic, random monoallelic gene expression in mammalian cells. Science, 343(6167):193–196, January 2014.

9. Mubeen Goolam, Antonio Scialdone, Sarah J L Graham, Iain C Macaulay, Agnieszka Jedrusik, Anna Hupalowska, Thierry Voet, John C Marioni, and Magdalena Zernicka-Goetz. Heterogeneity in oct4 and sox2 targets biases cell fate in 4-cell mouse embryos. Cell, 165(1):61–74, March 2016.

10. Diego Adhemar Jaitin, Ephraim Kenigsberg, Hadas Keren-Shaul, Naama Elefant, Franziska Paul, Irina Zaretsky, Alexander Mildner, Nadav Cohen, Steffen Jung, Amos Tanay, et al. Massively parallel single-cell rna-seq for marker-free decomposition of tissues into cell types. Science, 343(6172):776–779, 2014.

11. Diederik P. Kingma and Jimmy Ba. Adam: A method for stochastic optimization, 2017.

12. Vladimir Yu Kiselev, Tallulah S Andrews, and Martin Hemberg. Challenges in unsupervised clustering of single-cell rna-seq data. Nature Reviews Genetics, 20(5):273–282, 2019.

13. Vladimir Yu Kiselev, Kristina Kirschner, Michael T Schaub, Tallulah Andrews, Andrew Yiu, Tamir Chandra, Kedar N Natarajan, Wolf Reik, Mauricio Barahona, Anthony R Green, et al. Sc3: consensus clustering of single-cell rna-seq data. Nature methods, 14(5):483–486, 2017.

14. Aleksandra A Kolodziejczyk, Jong Kyoung Kim, Jason C H Tsang, Tomislav Ilicic, Johan Henriksson, Kedar N Natarajan, Alex C Tuck, Xuefei Gao, Marc Buhler, Pentao Liu, John C Marioni, and Sarah A Teichmann. Single cell RNA-sequencing of pluripotent states unlocks modular transcriptional variation. Cell Stem Cell, 17(4):471–485, October 2015.

15. T.K. Moon. The expectation-maximization algorithm. IEEE Signal Processing Magazine, 13(6):47–60, 1996.

16. Alex A Pollen, Tomasz J Nowakowski, Joe Shuga, Xiaohui Wang, Anne A Leyrat, Jan H Lui, Nianzhen Li, Lukasz Szpankowski, Brian Fowler, Peilin Chen, Naveen Ramalingam, Gang Sun, Myo Thu, Michael Norris, Ronald Lebofsky, Dominique Toppani, Darnell W Kemp, 2nd, Michael Wong, Barry Clerkson, Brittnee N Jones, Shiquan Wu, Lawrence Knutsson, Beatriz Alvarado, Jing Wang, Lesley S Weaver, Andrew P May, Robert C Jones, Marc A Unger, Arnold R Kriegstein, and Jay A A West. Low-coverage single-cell mRNA sequencing reveals cellular heterogeneity and activated signaling pathways in developing cerebral cortex. Nat. Biotechnol., 32(10):1053–1058, October 2014.

17. Peng Qiu. Embracing the dropouts in single-cell rna-seq analysis. Nature communications, 11(1):1169, 2020.

18. Peng Qiu. Embracing the dropouts in single-cell RNA-seq analysis. Nat. Commun., 11(1):1169, March 2020.

19. Francisca Rojas Ringeling, Stefan Canzar, et al. Linear-time cluster ensembles of large-scale single-cell rna-seq and multimodal data. Genome research, 31(4):677–688, 2021.

20. Karthik Shekhar, Sylvain W Lapan, Irene E Whitney, Nicholas M Tran, Evan Z Macosko, Monika Kowalczyk, Xian Adiconis, Joshua Z Levin, James Nemesh, Melissa Goldman, Steven A McCarroll, Constance L Cepko, Aviv Regev, and Joshua R Sanes. Comprehensive classification of retinal bipolar neurons by single-cell transcriptomics. Cell, 166(5):1308–1323.e30, August 2016.

21. Marlon Stoeckius, Christoph Hafemeister, William Stephenson, Brian Houck-Loomis, Pratip K Chattopadhyay, Harold Swerdlow, Rahul Satija, and Peter Smibert. Simultaneous epitope and transcriptome measurement in single cells. Nat. Methods, 14(9):865–868, September 2017.

22. Tabula Muris Consortium, Overall coordination, Logistical coordination, Organ collection and processing, Library preparation and sequencing, Computational data analysis, Cell type annotation, Writing group, Supplemental text writing group, and Principal investigators. Single-cell transcriptomics of 20 mouse organs creates a tabula muris. Nature, 562(7727):367–372, October 2018.

23. Vincent A Traag, Ludo Waltman, and Nees Jan Van Eck. From louvain to leiden: guaranteeing well-connected communities. Scientific reports, 9(1):5233, 2019.

24. Tianyu Wang, Boyang Li, Craig E Nelson, and Sheida Nabavi. Comparative analysis of differential gene expression analysis tools for single-cell rna sequencing data. BMC bioinformatics, 20(1):1–16, 2019.

25. Victor Wang, Pietro Antonio Cicalese, Anto Sam Crosslee Louis Sam Titus, and Chandra Mohan. Polaratio: A magnitude-contingent monotonic correlation metric and its improvements to scrna-seq clustering. bioRxiv, 2021.

26. Xin Wei, Ziyi Li, Hongkai Ji, and Hao Wu. EDClust: an EM–MM hybrid method for cell clustering in multiple-subject single-cell RNA sequencing. Bioinformatics, 38(10):2692–2699, 03 2022.

27. Liying Yan, Mingyu Yang, Hongshan Guo, Lu Yang, Jun Wu, Rong Li, Ping Liu, Ying Lian, Xiaoying Zheng, Jie Yan, Jin Huang, Ming Li, Xinglong Wu, Lu Wen, Kaiqin Lao, Ruiqiang Li, Jie Qiao, and Fuchou Tang. Single-cell RNA-Seq profiling of human preimplantation embryos and embryonic stem cells. Nat. Struct. Mol. Biol., 20(9):1131–1139, September 2013.

28. Matthew D Young, Thomas J Mitchell, Felipe A Vieira Braga, Maxine G B Tran, Benjamin J Stewart, John R Ferdinand, Grace Collord, Rachel A Botting, Dorin-Mirel Popescu, Kevin W Loudon, Roser Vento-Tormo, Emily Stephenson, Alex Cagan, Sarah J Farndon, Martin Del Castillo Velasco-Herrera, Charlotte Guzzo, Nathan Richoz, Lira Mamanova, Tevita Aho, James N Armitage, Antony C P Riddick, Imran Mushtaq, Stephen Farrell, Dyanne Rampling, James Nicholson, Andrew Filby, Johanna Burge, Steven Lisgo, Patrick H Maxwell, Susan Lindsay, Anne Y Warren, Grant D Stewart, Neil Sebire, Nicholas Coleman, Muzlifah Haniffa, Sarah A Teichmann, Menna Clatworthy, and Sam Behjati. Single-cell transcriptomes from human kidneys reveal the cellular identity of renal tumors. Science, 361(6402):594–599, August 2018.

29. Allen W Zhang, Ciara O’Flanagan, Elizabeth A Chavez, Jamie LP Lim, Nicholas Ceglia, Andrew McPherson, Matt Wiens, Pascale Walters, Tim Chan, Brittany Hewitson, et al. Probabilistic cell-type assignment of single-cell rna-seq for tumor microenvironment profiling. Nature methods, 16(10):1007–1015, 2019.

30. Martin Jinye Zhang, Angela Oliveira Pisco, Spyros Darmanis, and James Zou. Mouse aging cell atlas analysis reveals global and cell type-specific aging signatures. Elife, 10, April 2021.

31. Ze Zhang, Danni Luo, Xue Zhong, Jin Huk Choi, Yuanqing Ma, Stacy Wang, Elena Mahrt, Wei Guo, Eric W Stawiski, Zora Modrusan, et al. Scina: a semi-supervised subtyping algorithm of single cells and bulk samples. Genes, 10(7):531, 2019.

32. Grace X. Y. Zheng, Jessica M. Terry, Phillip Belgrader, Paul Ryvkin, Zachary W. Bent, Ryan Wilson, Solongo Batjargal Ziraldo, Tobias D. Wheeler, Geoffrey P. McDermott, Junjie Zhu, Mark T. Gregory, Joe Shuga, Luz Montesclaros, Jason G. Underwood, Donald A. Masquelier, Stefanie Y. Nishimura, Michael Schnall-Levin, Paul W Wyatt, Christopher M. Hindson, Rajiv Pranesh Bharadwaj, Alexander Wong, Kevin D Ness, Lan Beppu, H. Joachim Deeg, Christopher McFarland, Keith R. Loeb, Keith R. Loeb, W. J. Valente, W. J. Valente, Nolan G. Ericson, Emily A. Stevens, Jerald P. Radich, Tarjei Sigurd Mikkelsen, Benjamin J. Hindson, and Jason H. Bielas. Massively parallel digital transcriptional profiling of single cells. Nature Communications, 8, 2016.

